# A lysosomal lipid transport pathway that enables cell survival under choline limitation

**DOI:** 10.1101/2022.11.27.517422

**Authors:** Samantha G. Scharenberg, Wentao Dong, Kwamina Nyame, Roni Levin-Konigsberg, Aswini R. Krishnan, Eshaan S. Rawat, Kaitlyn Spees, Michael C. Bassik, Monther Abu-Remaileh

## Abstract

Lysosomes degrade macromolecules and recycle their nutrient content to support cell function and survival over a broad range of metabolic conditions. Yet, the machineries involved in lysosomal recycling of many essential nutrients remain to be discovered, with a notable example being choline, an essential metabolite liberated in large quantities within the lysosome via the degradation of choline-containing lipids. To identify critical lysosomal choline transport pathways, we engineered metabolic dependency on lysosome-derived choline in pancreatic cancer cells. We then exploited this dependency to perform an endolysosome-focused CRISPR-Cas9 negative selection screen for genes mediating lysosomal choline recycling. Our screen identified the orphan lysosomal transmembrane protein SPNS1, whose loss leads to neurodegeneration-like disease in animal models, as critical for cell survival under free choline limitation. We find that *SPNS1* loss leads to massive accumulation of lysophosphatidylcholine (LPC) and lysophosphatidylethanolamine (LPE) within the lysosome. Mechanistically, we revealed that SPNS1 is required for the efflux of LPC species from the lysosome to enable their reesterification into choline-containing phospholipids in the cytosol. Using cell-based lipid uptake assays, we determine that SPNS1 functions as a proton gradient-dependent transporter of LPC. Collectively, our work defines a novel lysosomal phospholipid salvage pathway that is required for cell survival under conditions of choline limitation, and more broadly, provides a robust platform to deorphan lysosomal gene functions.

## Introduction

Cells require a constant supply of nutrients to support their vital activities and growth. To survive inevitable fluctuations in nutrient availability, *eukaryotes* have evolved multiple strategies to acquire small molecules essential for their survival (*1*). Important among these are the endocytic pathways that enable internalization of nutrient-rich cargo that cannot translocate across the plasma membrane by passive diffusion or active transport. These pathways converge on the lysosome, which scavenges the vital nutrients from the microenvironment by degrading endocytosed cargos and liberating their constituent metabolites (*2*). Thus, nutrient recycling through the lysosome represents a critical source of metabolic adaptability for cells to survive various states of nutrient deprivation.

Despite their essential role in nutrient acquisition, the lysosomal machineries that mediate nutrient recycling, in particular the transport of liberated nutrients to the cytosol, are not fully defined. In recent work, we demonstrated that the lysosomal transporter SLC38A9 becomes selectively essential for cell growth under leucine-scarce conditions, while it is dispensable under leucine-replete conditions, leading to the discovery that SLC38A9 functions to efflux leucine derived from lysosome-degraded proteins (*3*). This work illustrates that metabolic dependency on lysosomes engineered through depleting culture media of specific free nutrients can be exploited to discover lysosomal proteins involved in recycling these nutrients.

One essential nutrient recycled through the lysosome is choline (*4*, *5*). Choline is a small, cationic metabolite that constitutes the polar head group of the most abundant cellular phospholipid (phosphatidylcholine, PC; comprising >50% of cellular membranes) and sphingolipid (sphingomyelin, SM). It is also required for the synthesis of the neurotransmitter acetylcholine and serves as a methyl-donor in one-carbon metabolism (*6*). Through the degradation of choline-containing lipids, large quantities of choline-containing catabolites are produced and exported from the lysosome to be recycled in the cytosol (**Fig. 1a**) (*7*, *8*). Despite their high flux through the lysosome, many questions remain about the identity of the choline-containing compounds being recycled and the machineries mediating this process.

**Figure 1:**
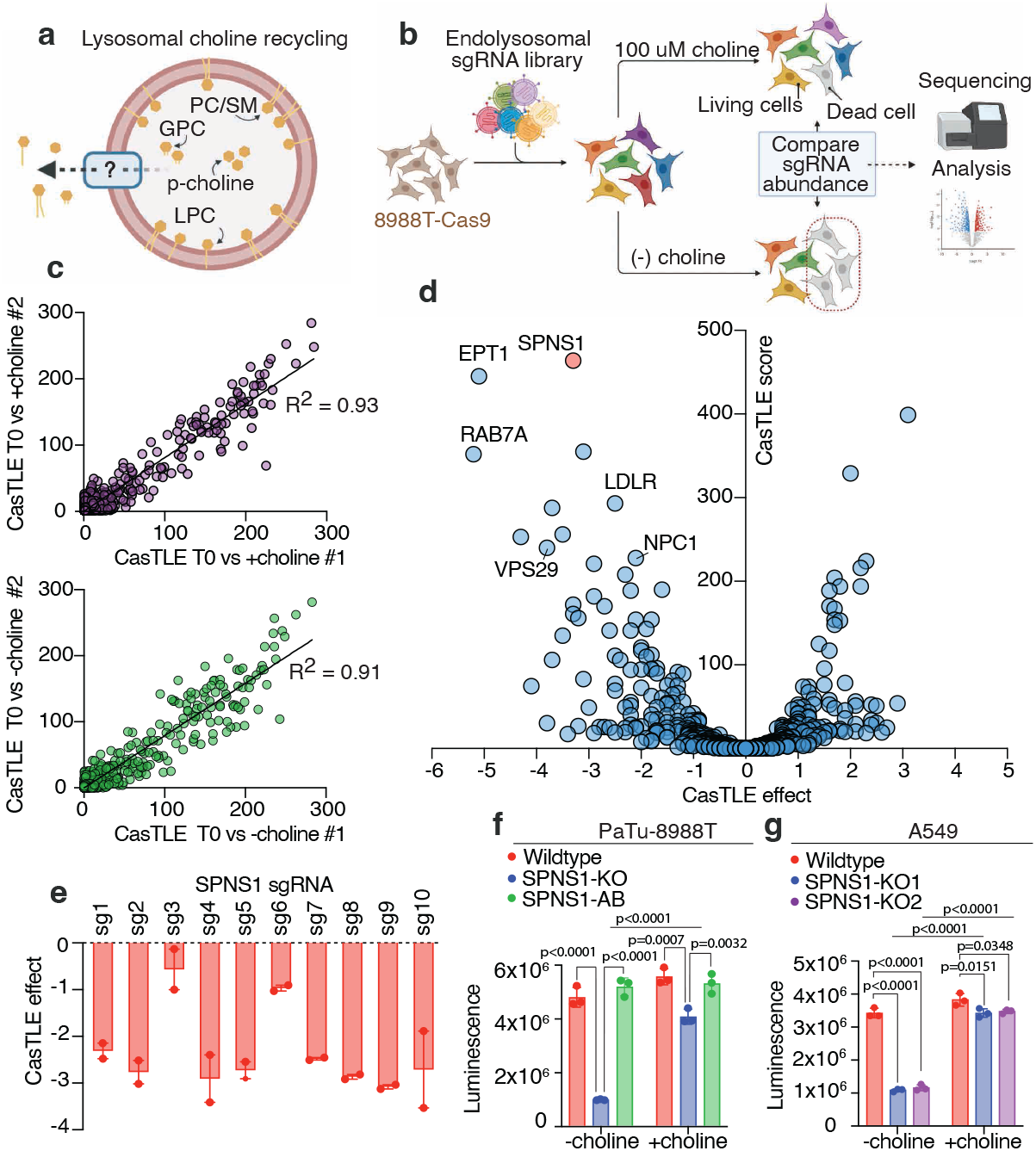
Endolysosomal CRISPR-Cas9 screen implicates SPNS1 in the response to choline deprivation. **a)** Overview of lysosomal choline recycling highlighting an unknown transporter that exports choline-containing catabolites. PC: phosphatidylcholine, SM: sphingomyelin, GPC: glycerophosphocholine, p-choline: phosphocholine, LPC: lysophosphatidylcholine. **b)** Schematic overview of the endolysosomal CRISPR-Cas9 negative selection screen performed to identify genes required for lysosomal choline recycling. **c)** Correlation plots between biological replicates in the endolyososmal screen for each culture condition (top: +choline condition; bottom: -choline condition). Plots depict CasTLE scores computed between time 0 and fourteen doublings for each replicate in the indicated culture condition. Simple linear regression was performed with R^2^ values indicated on the plots. **d)** Volcano plot for -choline vs +choline CasTLE score vs the effect for each gene in the endolysosomal library (CasTLE effect). Endolysosomal genes that are essential in the -choline condition have a negative CasTLE effect size. The highest-scoring gene (SPNS1) is highlighted in red. Tabulated data are found in Supplementary table 2. **e)** Individual CasTLE effects for -choline vs. +choline for all SPNS1-targeting sgRNAs in the endolysosomal library. Data show mean +/− SEM. **f)** Live cell abundance measured by CellTiter-Glo of wildtype, SPNS1-KO, and SPNS1-AB PaTu-8988T cells after five days in -choline and +choline cultures. Data show mean +/− SEM. Statistical test: one-way ANOVA with Tukey’s HSD post-hoc. **g)** Live cell abundance measured by CellTiter-Glo of wildtype, SPNS1-KO1, and SPNS1-KO2 A549 cells after five days in -choline and +choline cultures. Data show mean +/− SEM. Statistical test: one-way ANOVA with Tukey’s HSD post-hoc.

Here, we develop and apply a customized CRISPR-Cas9 screen coupled with engineered choline-depleted culture medium to identify a novel choline-phospholipid salvage pathway orchestrated by the lysosome. We find that lysosomes enable cell survival under choline deprivation by supplying lysophospholipids, in particular lysophosphatidylcholine (LPC), to augment phospholipid production in the cytoplasm. This novel mechanism is dependent on the orphan lysosomal major facilitator superfamily transporter *SPNS1*, the loss of which leads to substance storage and neurodegeneration-like disease in animal models (*6*, *9*–*12*). We demonstrate that SPNS1 functions to export LPC species from the lysosomal lumen to the cytosol for re-acylation to many PC species, and we posit that by supplying choline-containing scaffolds for PC synthesis, lysosomal LPCs compensate for loss of exogenous choline. This work thus reveals a novel pathway for lysophospholipid export from the lysosome that supports synthesis of vital phospholipids, and more broadly, establishes a generalizable approach for deorphanizing lysosomal genes and pathways.

## Results

### A forward genetics approach reveals *SPNS1* as an essential lysosomal gene under choline-limited conditions

We speculated that growing cells in media depleted of free choline would force their dependence on lysosomal recycling of choline, thereby conferring essentiality to lysosomal choline-scavenging genes. This engineered dependency, coupled with high-throughput negative-selection genetic screening, might represent an efficient, unbiased approach to identify novel lysosomal genes involved in recycling choline-containing molecules.

To this end, we designed a CRISPR-Cas9 library targeting all endolysosomal genes as well as a selective subset of metabolic and signaling genes (1,061 genes with 10 sgRNAs per gene, endolysosomal library *hereafter*) (**Supplementary Table 1** and methods). Using this library, we sought to determine endolysosomal genes involved in choline recycling by screening for those that become essential under choline deprivation (**Fig. 1a&b**). We selected the pancreatic cancer cell line PaTu-8988T for screening because these cells survive choline deprivation while still exhibiting modest sensitivity to choline restriction (**Fig. S1a&b**), indicating an active choline scavenging pathway. This behavior is consistent with the pancreatic ductal adenocarcinoma (PDAC) origin of PaTu-8988T cells, a cancer known for adapting to nutrient scarce microenvironment (*13*–*16*).

PaTu-8988T cells expressing Cas9 were infected with the endolysosomal library and propagated for 14 doublings in culture media lacking free choline (-choline) or supplemented with 100 μM choline (+choline) (**Fig. 1b**). A high correlation between replicates in both the -choline and +choline conditions was observed (**Fig. 1c**). Gene scores and depletion/enrichment effects between the -choline and +choline conditions were determined using CasTLE analysis (*17*), revealing genes whose targeting sgRNAs are significantly enriched or depleted in the -choline condition (**Fig. 1d** and **Supplementary Table 2**). Genes whose targeting sgRNAs are depleted in the -choline condition are considered essential for cells to grow in the absence of choline. Among the highly scoring genes in this category is *EPT1,* a phosphotransferase that catalyzes the synthesis of the phospholipid phosphatidylethanolamine (PE), a direct lipid precursor of PC (*18*). PE is trimethylated to PC by the enzyme PE N-methyltransferase (encoded for by *PEMT)* and serves as an important alternative substrate for PC synthesis in the absence of free choline (*6*); moreover, PC synthesized from PE can be catabolized to produce free choline. Thus, *EPT1* represents a positive control in this screen (*18*–*20*) (**Fig. 1d**).

In addition, a cluster of genes necessary for trafficking macromolecules to the lysosome were identified to be required for cell growth in the -choline condition (**Fig. 1d**). Of these are *RAB7A,* encoding a small GTPase responsible for regulating late endocytic transport to the lysosome (*21*, *22*); *LDLR,* a plasma membrane receptor that mediates the uptake and delivery of lipid-rich extracellular lipoproteins to the lysosome (*23*, *24*); and *VPS29,* encoding a retromer complex component necessary for recycling hundreds of transmembrane proteins, including LDLR, from degradation pathways back to the cell surface for continuous cargo trafficking to the lysosome (*25*). Altogether, these results indicate that faithful trafficking of choline-containing cargo (mainly phospholipids) to lysosomes is essential for survival under choline-restricted conditions, supporting an essential role for lysosome-mediated phospholipid catabolism in compensating for choline restriction.

To identify the elusive lysosomal choline-containing catabolite transporter, we focused on transmembrane proteins. Interestingly, *NPC1,* the lysosomal permease for cholesterol and, putatively, sphingosine (*26*–*28*) became more essential in the -choline condition (**Fig. 1d, Fig. S1c**). NPC1 loss causes Niemann–Pick type C, a fatal autosomal-recessive lipid-storage disorder characterized by progressive neurodegeneration (*29*). NPC1 deficiency is known to disrupt sphingomyelin catabolism (*27*), a potential source of choline, which might explain its essentiality in the -choline condition. Notably, the top-scoring gene whose targeting guides were depleted in the -choline condition was *SPNS1* (**Fig. 1d**), which encodes SPNS1, a lysosomal transmembrane protein and a member of the major facilitator superfamily (MFS) previously hypothesized to export sugars (*9*) or sphingolipids (*30*). Deficiency in *SPNS1* homologs results in neurodegeneration and lysosomal storage disease-like presentation in model organisms (*9*–*12*), mirroring biochemical pathologies associated with deficiency of lysosomal solute carriers. All ten sgRNAs targeting *SPNS1* were more depleted in the -choline compared to +choline condition, indicating the robustness of the *SPNS1*-depletion phenotype under screening conditions (**Fig. 1e**).

To validate that *SPNS1* is an essential gene during free choline restriction, we generated *SPNS1*-knockout (KO) cells in two cell lines: PaTu-8988T pancreatic cancer cells and A549 lung adenocarcinoma cells (**Fig. S1d&e**). We compared growth between *SPNS1-*KO cells and wildtype cells cultured in -choline and +choline conditions for five days. Consistent with our screen results, we observed a dramatic growth defect in *SPNS1*-KO cells compared to wildtype cells in -choline, compared to only minor defect in +choline conditions for both PaTu-8988T and A549 lines (**Fig. 1f&g**). Importantly, re-expressing *SPNS1* in *SPNS1*-KO cells (*SPNS1*-addback, *hereafter SPNS1*-AB) completely rescued cell growth defects (**Fig. 1f and Fig. S1f**). Together, these results indicate that SPNS1 function is required for cell growth under -choline conditions.

### Loss of *SPNS1* leads to cell death under choline deprivation

To further characterize how loss of *SPNS1* affects cell growth, we generated growth curves for PaTu-8988T *SPNS1-KO* cells in -choline and +choline conditions and compared these to the corresponding growth curves of wildtype and *SPNS1-AB* cells. To our surprise, cell growth was initially similar across the three different genotypes in +choline and -choline cultures (**Fig. 2a**). However, around three to four days in -choline medium, *SPNS1*-KO cell number progressively declined while the wildtype and *SPNS1-AB* cells continued to increase in number. By day six in -choline medium, there was a significant decrease in the number of viable *SPNS1*-KO cells compared to wildtype and *SPNS1*-AB cells in the -choline condition (**Fig. 2a**). Decline in viable cell count was not observed in the +choline condition (**Fig. 2a**).

**Figure 2:**
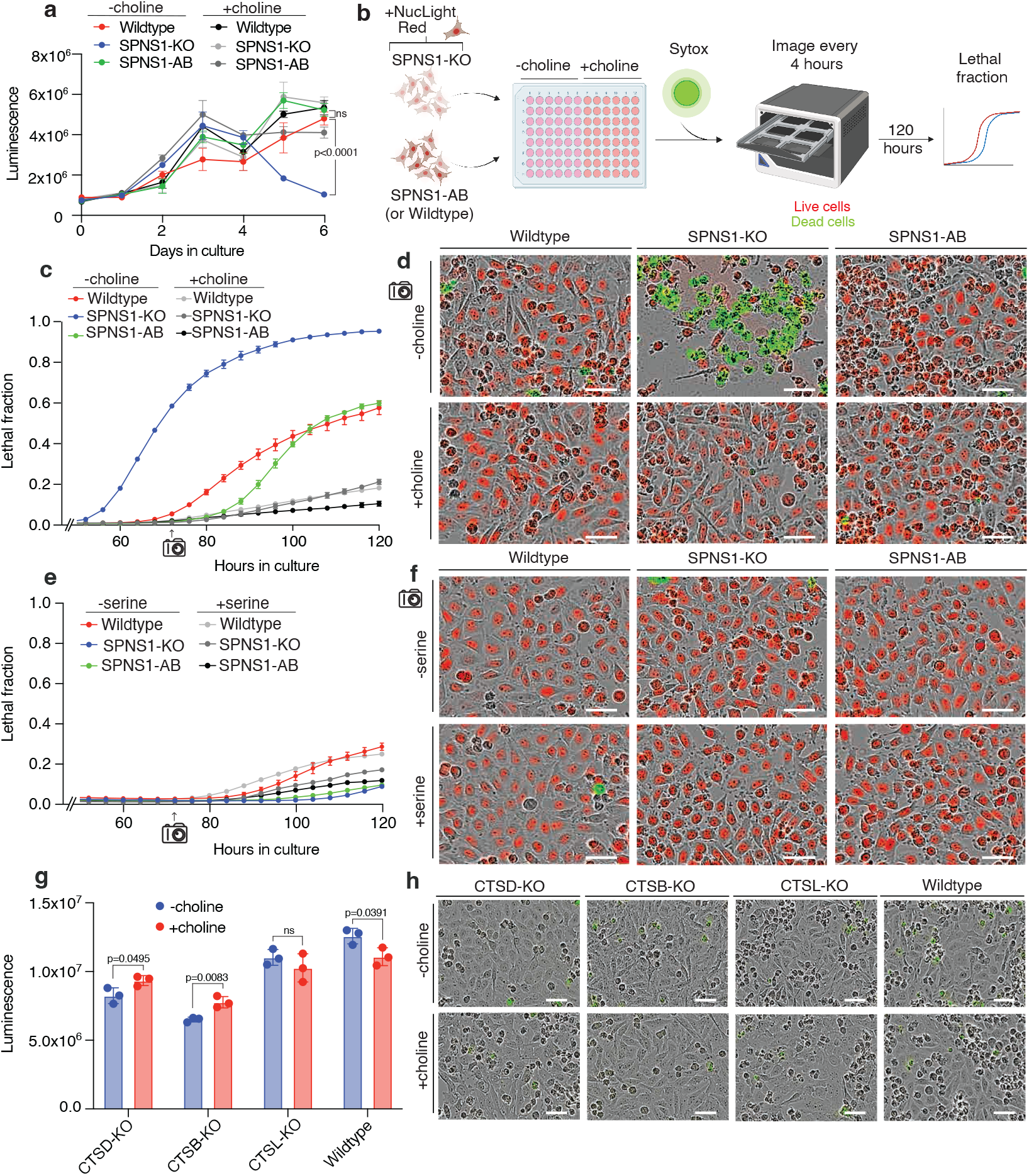
SPNS1 deficiency promotes cell death under choline-limiting conditions. **a)** Cell growth curves generated for PaTu-8988T wildtype, SPNS1-KO, and SPNS1-AB, cells in choline-depleted medium (-choline) or medium supplemented with 100 μM choline (+choline). Data show mean +/− SEM. Statistical test: two-tailed unpaired t-test performed individually at each time point indicated. **b)** Schematic for Incucyte workflow used to generate lethal fraction curves. **c)** Lethal fraction of wildtype, SPNS1-KO, and SPNS1-AB cells grown in choline-depleted medium (-choline) or medium supplemented with 100 μM choline chloride (+choline). Data show mean +/− SEM. **d)** Representative Incucyte fields from (c) at the 72-hour time-point. Green marks dead cells stained with Sytox Green; red marks nuclei of live cells. White scale bar represents 50 μm. **e)** Lethal fraction of wildtype, SPNS1-KO, and SPNS1-AB cells grown in serine-depleted medium (-serine) or medium supplemented with 27 mg/L serine (+serine). Data show mean +/− SEM. **f)** Representative Incucyte fields from (e) at the 72-hour time-point. Green marks dead cells stained with Sytox Green; red marks nuclei of live cells. White scale bar represents 50 μm. **g)** Live cell abundance of PaTu-8988T knockouts for Cathepsin D (CTSD), Cathepsin B (CTSB) and Cathepsin L (CTSL), and wildtype cells after culturing for five days in choline-depleted medium (-choline) or medium supplemented with 100 μM choline chloride (+choline). Data show mean +/− SEM. Statistical test: two-tailed unpaired t-test. **h)** Representative images of cells from (g) stained with propidium iodide (PI) for detection of dead cells (PI is pseudo-colored to green for clarity and consistency with dead cell representation in earlier panels. White scale bar represents 50 μm.

To verify that the decline in viable *SPNS1-KO* cell count resulted from cell death, we used the Incucyte system to monitor the lethal fraction of *SPNS1*-KO, wildtype and *SPNS1-AB* cells over five days in +choline and -choline conditions (*31*) (**Fig. 2b**). Consistent with the growth curves, *SPNS1*-KO cells in the -choline, but not +choline, condition exhibited a remarkable, rapid loss of viability at approximately three days in culture (**Fig. 2c&d, Fig. S2a&b**). The cell death exhibited by *SPNS1*-KO cells preceded any loss in cell viability of wildtype cells in -choline by approximately twenty-four hours (**Fig. 2c&d, Fig. S2a&b**) and was observed in a second PaTu-8988T *SPNS1*-KO cell line (**Fig S2c&d**), supporting the reproducibility of the phenotype. Rapid cell viability loss was not observed in the +choline condition for any genotype (**Fig. 2c&d, Fig. S2b**). Notably, re-expressing *SPNS1* in *SPNS1-KO* cells rescued their ability to survive choline deprivation to wildtype levels (**Fig. 2c&d, Fig. S2a&b**).

Adaptation to metabolic stress is a major lysosomal function (*32*, *33*), and loss of *SPNS1* might have caused a general dysfunction that limited the ability of lysosomes to supply various nutrients including choline. To test that loss of cell viability in *SPNS1-KO* cells is specific to choline depletion, we generated lethal fraction curves for *SPNS1-*KO, wildtype and *SPNS1-AB* PaTu-89898T cells in media lacking a different cellular nutrient that is similarly a constituent of polar lipid headgroups: serine. While cell growth was slowed in the -serine condition across all genotypes, no *SPNS1*-dependent loss in cell viability was observed in -serine compared to +serine conditions (**Fig. 2e&f and Fig. S2e&f**). To further establish that the death of *SPNS1-KO* cells in -choline condition is due to a specific interaction between choline and *SPNS1* function, we generated PaTu-8988T cells deficient for the lysosomal proteases CSTD, CTSB and CTSL, whose loss causes lysosomal dysfunction and human diseases (*29*, *34*–*36*). Consistent with their role in maintaining lysosomal homeostasis, loss of these proteases leads to a general growth defect in +choline and -choline conditions compared to wildtype cells, with cells in -choline medium being slightly more affected in some cases (**Fig. 2g**). However, we did not observe any cell death in *CSTD, CTSB* or *CTSL* KO cells growing in -choline medium (**Fig. 2h**). These results establish that SPNS1 supports cell survival under choline limitation through a function that is related to choline scavenging.

### SPNS1-deficient lysosomes accumulate lysophospholipids

Based on these results demonstrating that *SPNS1* is essential under choline restriction, we reasoned that SPNS1 is required for lysosomal choline recycling. Furthermore, because SPNS1 is localized to the limiting lysosomal membrane and is structurally homologous to solute carriers in the major facilitator superfamily (*37*), we initially hypothesized that SPNS1 is a transporter of either choline or phosphocholine, as these would directly contribute to choline pools in the cytosol, and their efflux from the lysosome has been previously reported (*7*, *8*).

The substrate of a lysosomal transporter mediating efflux would be predicted to accumulate in the lysosome upon loss of the transporter, as is observed in many human lysosomal storage diseases (*34*). Intralysosomal substrate accumulation can be robustly quantified using our rapid lysosomal immunopurification (LysoIP) coupled with mass spectrometry (*3*, *4*, *38*). We first adapted and optimized our LysoIP pipeline to PaTu-8988T cells (**Fig. 3a, Fig. S3a** and methods). We demonstrated efficient pull-down of PaTu-8988T lysosomes via capture of both lysosomal transmembrane and intraluminal proteins (**Fig. 3b**), as well as intralysosomal markers and metabolites (**Fig. 3c, Fig. S3b**).

**Figure 3:**
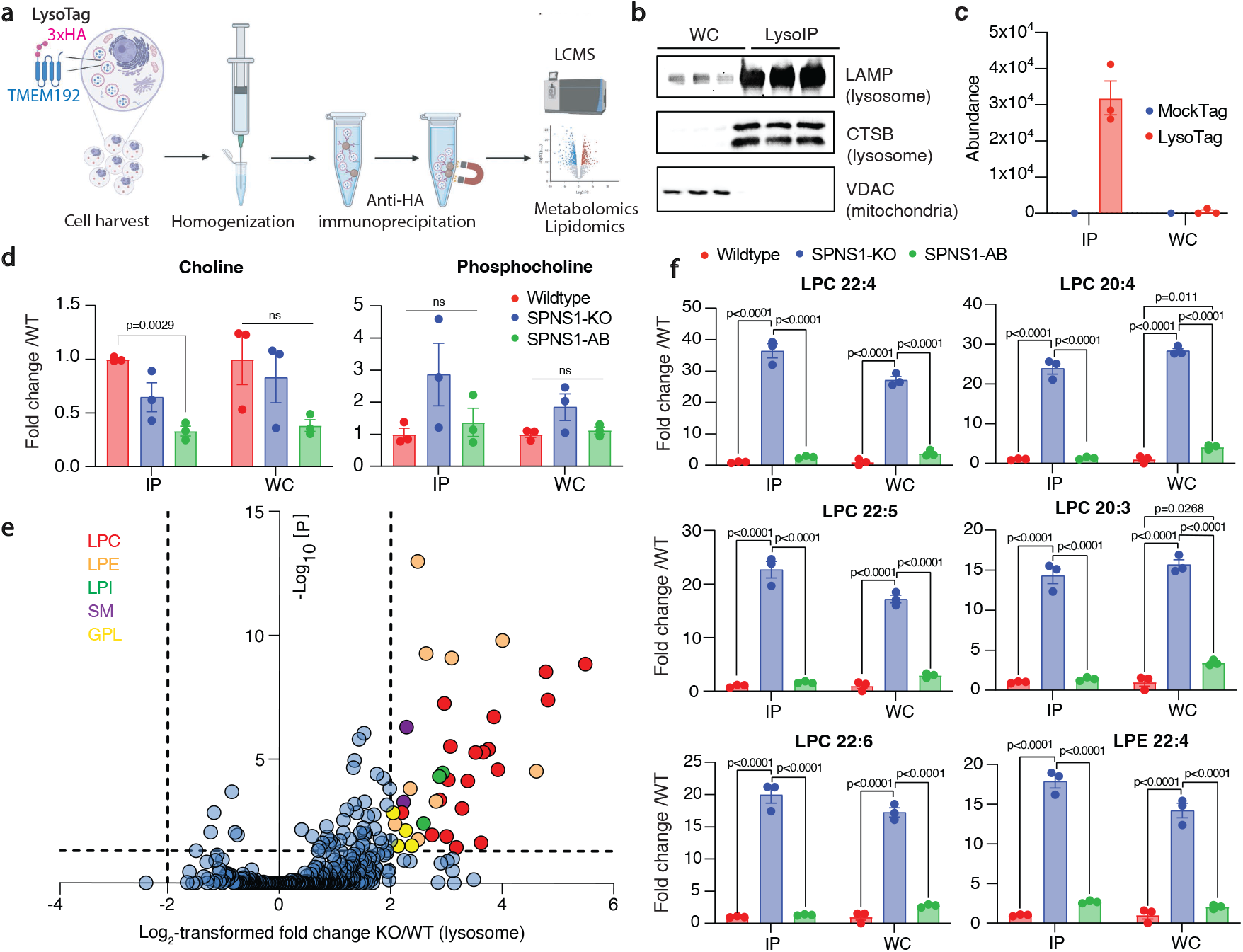
SPNS1-deficient lysosomes accumulate LPCs and LPEs. **a)** Schematic of LysoIP workflow used in this study. **b)** Immunoblot analysis for LysoIP validation on wildtype PaTu-8988T cells expressing LysoTag. Blot depicts three biological replicates. **c)** Lysotracker quantification in LysoIP (IP) and whole-cell (WC) fractions from wildtype PaTu-8988T cells expressing LysoTag (TMEM192-3xHA; n=3; Data show mean +/− SEM.) versus MockTag (TMEM192-3xflag; n=1). **d)** Quantitation of lysosomal (IP) and whole-cell (WC) abundance of choline and phosphocholine (normalized to a pool of endogenous amino acids: phenylalanine, methionine and tyrosine in positive ion mode) in wildtype, SPNS1-KO and SPNS1-AB cells. Data show mean of fold change versus wildtype +/− SEM. Statistical test: one-way ANOVA with Tukey’s HSD post-hoc. **e)** Volcano plot presentation of untargeted lipidomics data (SPNS1-KO (KO) vs wildtype (WT)) highlighting lysophosphatidylcholines (LPC; red), lysophosphatidylethanolamines (LPE; orange), lysophosphatidylinositol (LPI; green), sphingomyelin (SM; purple) and glycerophospholipids (GPL; yellow). Oneway ANOVA with post-hoc Tukey HSD test was performed for P-values. Adjust P-values were calculated by Benjamini–Hochberg correction for the false-discovery rate at 5%. **f)** Quantitation of the top six accumulated LPCs and LPEs (normalized to internal standards from SPLASH LIPIDOMIX) in lysosomes (IP) and whole-cell (WC) fractions of wildtype, SPNS1-KO and SPNS1-AB cells. Data show mean of fold change versus wildtype +/− SEM. Statistical test: one-way ANOVA with Tukey’s HSD post-hoc.

Surprisingly, we observed no increase in the lysosomal levels of choline or phosphocholine upon *SPNS1* loss, but a minor decrease if any (**Fig. 3d, Fig. S3c**). However, in the lysosomal metabolome derived from *SPNS1*-KO cells, we noticed striking elevations in features annotated as choline and phosphocholine based on *m/z* but elute at early retention times corresponding to lipids (**Fig. S3d**). We predicted that these must be the result of in-source fragmentation of choline- and phosphocholine-containing lipids. Indeed, by re-analyzing our polar metabolite data to include lysophospholipids, we found massive accumulation of lipid catabolites tentatively annotated as LPCs and lysophosphatidylethanolamines (LPEs) in *SPNS1*-KO lysosomes (**Fig. S3e**). These lysophospholipids are generated by the hydrolysis of one acyl group of their corresponding phospholipid. Of importance, the high abundance of LPCs and the matching retention time to that of the observed in-source fragments of choline and phosphocholine indicate that they are the source of detected features (**Fig. S3d**).

While polar metabolite analysis can tentatively identify lysophospholipids, lipid analysis is needed to confirm their identity (*39*). We therefore performed unbiased lysosomal lipidomics and confirmed that LPCs and LPEs accumulate in lysosomes derived from *SPNS1-KO* cells (**Fig. 3e&f and Supplementary table 3**), with conclusive molecular evidence at the tandem mass spectrometry level (**Fig. S3f**). Targeted analyses further validated these results and demonstrated that re-expressing *SPNS1* completely rescued lysophospholipid accumulation (**Fig. 3f**). Importantly, due to the magnitude of the effect size in the lysosome, this accumulation is also observed at the whole-cell level (**Fig. 3f**). This result indicates that lysophospholipids are maintained at low levels in the cell, and an increase in their levels in the lysosome, despite its relatively small volume, can be detected in whole-cell metabolomics. Lastly, both untargeted lipidomics (**Fig. 3e**) and targeted quantitation (**Fig. S3g**) show that the level of sphingosine is not affected by the presence of *SPNS1.* This suggests that SPNS1 does not function as a sphingosine transporter as previously predicted (*30*). Altogether, our data support that SPNS1 is required for the clearance of LPCs and LPEs from the lysosome.

### SPNS1 exports LPCs from lysosomes for recycling to PC in the cytosol

Based on our observations, we proposed that SPNS1 is a lysosomal lysophospholipid transporter. SPNS1 belongs to the major facilitator superfamily of transporters, though its transported substrate has not yet been identified. The missense point mutation E217 in the *Drosophila melanogaster* homolog *Spinster* was discovered as a loss-of-function mutant causing a lysosome storage disease-like phenotype (*10*). Multiple sequence alignments of human SPNS1 and Spinster suggest E217 to be a conserved residue corresponding to E164 in human SPNS1 (**Fig. 4a**). We speculated that E164 is either close to or within a pocket that mediates binding to potential substrate. We docked the structure of LPC (22:4) onto the predicted human SPNS1 structure from I-TASSER using Schrödinger Glide and identified E164 to be in close proximity with the docked LPC substrate, suggesting its possible involvement in stabilizing LPC during transport (**Fig. 4b**). Additional conserved residues, including R76 and H427, were found to potentially assist in LPC stabilization (**Fig. 4b**). Our *in silico* modeling supports the assumption that the structure of SPNS1’s is compatible with its putative function as a lysophospholipid transporter.

**Figure 4:**
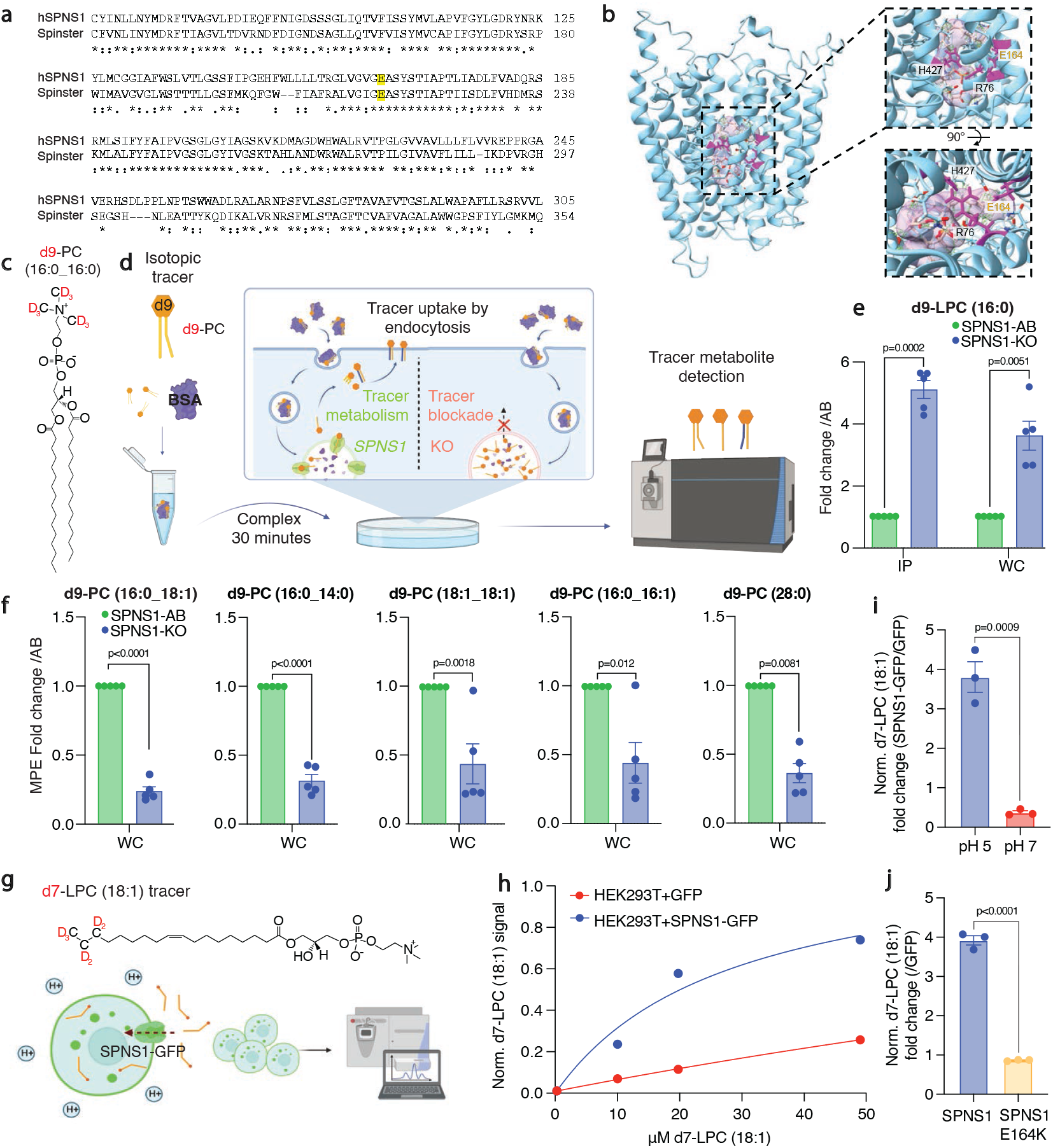
SPNS1 is the lysosomal transporter of LPC. **a)** Multiple sequence alignment between human SPNS1 and drosophila homolog spinster indicates the presence of fully conserved residues (marked with an asterisks). The conserved glutamic acid residue responsible for loss-of-function mutations in spinster, E217, corresponds to E164 in human SPNS1 (is highlighted in yellow). **b)** Schrödinger Glide-based docking of LPC (22:4) onto to the predicted human SPNS1 I-TASSER structure indicates that conserved residues, including E164, may be involved in LPC (22:4) stabilization. **c)** Structure of d9-PC (16:0) tracer used in this study. **d)** Schematic of d9-PC (16:0) tracing assay used in e and f. **e)** Quantitation of d9-LPC (16:0) in the lysosomal and whole-cell fractions of SPNS1-KO and SPNS1-AB cells expressed as paired-sample fold-change. Data show mean +/− SEM. Statistical test: two-tailed paired t-test. **f)** Quantitation of molar percent enrichment (MPE) for selected phosphatidylcholines expressed as paired-sample fold-change. Data show mean +/− SEM. Statistical test: two-tailed paired t-test. **g)** Schematic of d7-LPC (18:1) surface transport assay and d7-LPC (18:1) structure. **h)** Dose-response curve for d7-LPC (18:1) transport in HEK293T cells over-expressing GFP or over-expressing SPNS1-GFP. Data show d7-LPC (18:0) normalized to the endogenous lipid POPC. **i)** d7-LPC (18:1) transport in HEK293T cells overexpressing GFP or SPNS1-GFP at acidic pH (pH 5) and neutral pH (pH 7). Data are represented as fold change of d7-LPC (18:1) signal in HEK293T+SPNS1-GFP over HEK293T+GFP. Data show mean +/− SEM. Statistical test: two-tailed unpaired t-test. **j)** d7-LPC (18:1) transport in HEK293T cells expressing SPNS1-GFP or SPNS1(E164K)-GFP (expressed as fold change over HEK293T+GFP). The assay was performed with 20 μM d7-LPC (18:1) at pH 5. Data show mean +/− SEM. Statistical test: two-tailed unpaired t-test.

To experimentally validate SPNS1 as a transporter of lysophospholipids, we focused on LPC due to 1) its direct involvement in choline metabolism, and 2) its extreme accumulation in *SPNS1*-KO lysosomes. First, we leveraged isotope tracing (as previously described: (*38*, *40*)) to assess if LPC generated from lysosome-targeted deuterated PC is trapped in the lysosomes of *SPNS1*-deficient cells (**Fig. 4c&d**). After verifying equal tracer uptake by *SPNS1*-KO and *SPSN1*-AB cells (**Fig S4a**), we found a robust 5-6 fold accumulation of deuterated LPC in *SPNS1*-deficient lysosomes, supporting the putative role of SPNS1 in lysophospholipid efflux (**Fig. 4e**). Second, we assessed the abundance of labeled PCs in the whole-cell fractions of *SPNS1*-KO and *SPNS1*-AB cell, as LPCs transported from lysosomes are expected to contribute to PC pools in the cytoplasm upon re-esterification via the Lands Cycle (*41*, *42*). Using whole-cell isotope-tracing, we found that SPNS1 is required for LPC generated in the lysosome to contribute to PC pools in the cytoplasm, presumably by enabling LPC egress from lysosomes (**Fig. 4f**). Of importance to this assumption, loss of SPNS1 did not reduce basal choline phospholipid biosynthesis or turnover, as similar or higher labeling of PC species was observed in *SPNS1*-KO compared to *SPNS1*-AB cells when free d9 choline was used as a tracer (**Fig S4b**).

Finally, to provide definitive evidence for the role of SPNS1 in LPC transport, we leveraged a high exogenous expression system to over-express *SPNS1*-GFP in HEK293T cells, leading to spilling-over of *SPNS1*-GFP into the plasma membrane (**Fig. S4c**). Using a cell-based uptake assay with an *in vitro* proton gradient out-to-inside the cell (**Fig 4g**), we found that *SPNS1*-GFP mediates a robust transport of LPC across the membrane (**Fig. 4h**) in a pH-dependent manner (**Fig. 4i**), consistent with its role as lysosomal transporter of lysophospholipids. Mutating the conserved glutamate residue at position 164 to a lysine to generate *SPNS1*-E164K abolished transport activity to background levels (**Fig. 4j and Fig. S4c**). Taken together, our *in silico* modeling, lysosome lipid isotope tracing, and cellular transport assays indicate that SPNS1 transports LPCs from lysosomes to be used in phospholipid anabolism through a novel salvage pathway that is essential under choline limitation.

## Discussion

Our work identifies the orphan transmembrane protein SPNS1 as a transporter for LPC (and putatively, LPE) that is necessary for lysosomal phospholipid salvage and recycling. We demonstrate that SPNS1-derived LPCs support phospholipid synthesis under basal conditions, and notably, become essential under choline restriction when free choline for *de novo* synthesis of phospholipids is limited (**Fig. 5**).

**Figure 5:**
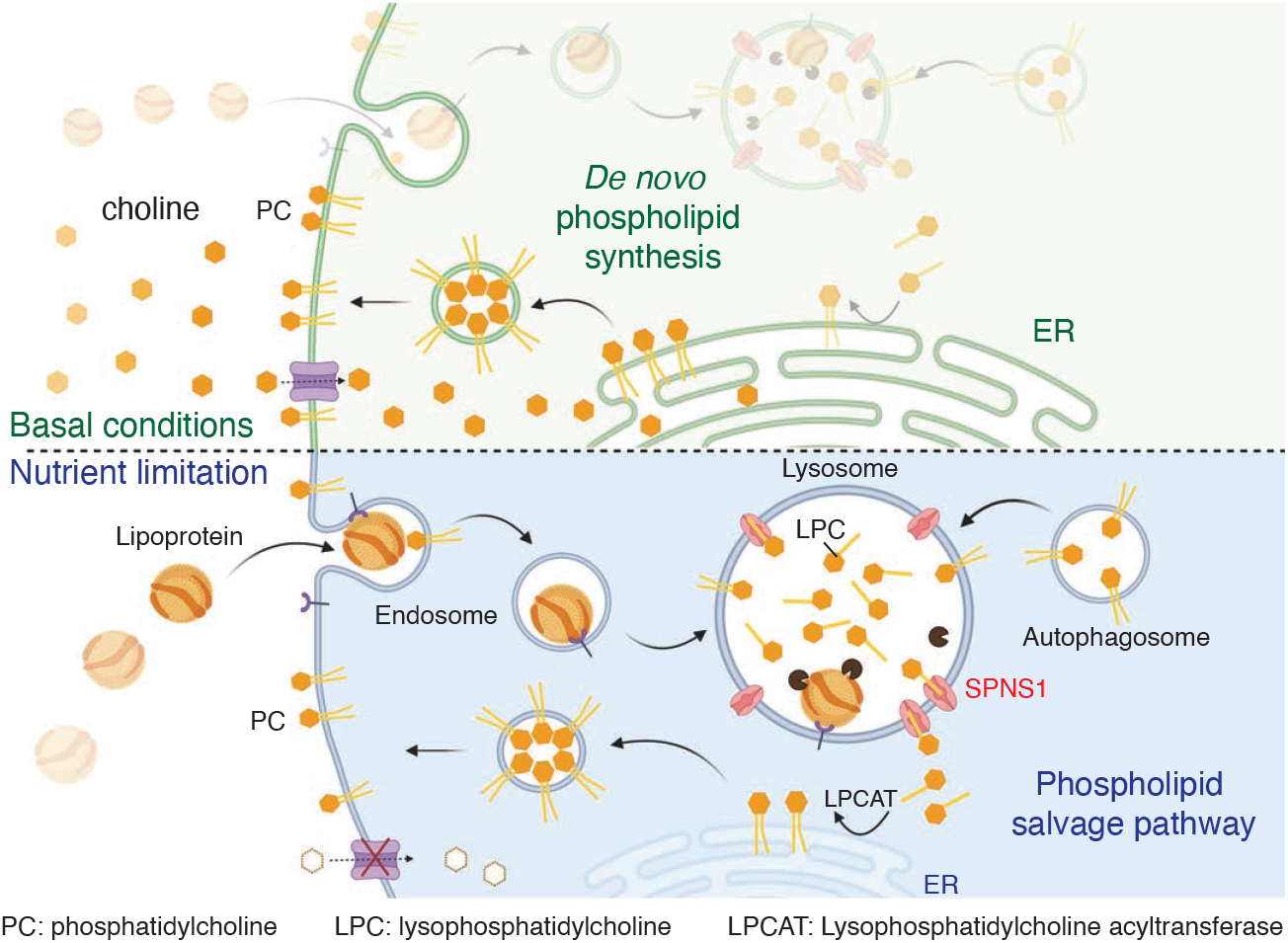
SPNS1-mediated novel lysosomal phospholipid salvage pathway. Model for a lysosomal lipid salvage pathway in which SPNS1 transports lysophosphatidylcholine to the cytosol for re-acylation to phosphatidylcholine. The pathway becomes essential for cell survival under choline-limited conditions when substrates for de novo phosphatidylcholine synthesis are restricted.

Biochemical studies have previously established that lysosomes recycle choline-related catabolites derived from the degradation of PC and SM (*7*, *8*). Seminal experiments by De Duve demonstrated that rat liver lysosomal extracts hydrolyze phospholipids to liberate glycerophosphodiesters (GPDs), including glycerophosphocholine (GPC), as terminal catabolites (*7*). Our group recently discovered that GPDs are exported from the lysosome in a CLN3-dependent manner (*38*). Similarly, early work using radiolabeled choline tracers revealed that SM is hydrolyzed to ceramide and phosphocholine by lysosomal acid sphingomyelinase to produce soluble phosphocholine, which exits the lysosome through a yet unidentified transporter (*8*). Once GPC and phosphocholine reach the cytosol, they enter pathways for incorporation into newly synthesized PCs and SMs (*5*, *8*, *38*). The biochemical mechanism proposed to explain these observations has been that GPDs and phosphocholine are the terminal choline-containing catabolites exported from the lysosome (*7*).

While we initially set out to identify the elusive phosphocholine transporter, our data provide strong evidence that an active machinery to transport LPC, and potentially its close lipid LPE, from lysosomes also exists. LPC and LPE are formed by the mono-deacylation of PC and PE, respectively, and are themselves deacylated once more to produce the di-deacylated catabolites GPC and GPE, respectively (*7*). Lysophospholipid export from the lysosome therefore prevents phospholipid catabolism to the terminal GPC and GPE catabolites, salvaging lysophospholipid intermediates for direct reacylation to various PC species – a stratagem that is easily rationalized as conserving energy equivalents that would otherwise be consumed in a futile deacylation-reacylation cycle between LPC/GPC and LPE/GPE. In support of this model, our isotope tracing establishes that salvaged LPC is re-acylated to PC upon export to the cytosol, likely via a one-step re-esterification mechanism utilizing ER-resident enzyme lysophosphatidylcholine acyltransferase (LPCAT) (*41*). Importantly, lysophospholipid salvage is a mechanism that can sustain phospholipid synthesis under conditions of critical nutrient limitation. Consistent with this, we establish that the machinery for LPC transport, namely SPNS1, is essential under choline limitation.

It is additionally plausible that SPNS1-supplied lysosomal LPCs have a dedicated cellular function. Studies have implicated *SPNS1* in autophagosome formation (*43*), a process requiring PC synthesis for membrane nucleation (*44*). Depletion of *SPNS1* protein leads to defective autophagic lysosome reformation, ultimately affecting autophagy-dependent processes such as synaptic pruning and neurodevelopment (*45*). Hence, it is possible that SPNS1-supplied lysosomal LPCs are shunted toward biosynthesis of PC specifically for autophagosome membranes. Lysophospholipids are also themselves functional, bioactive signaling molecules that mediate a variety of cellular responses via specific G-protein coupled receptors (GPCR) (*46*). Lysosomal LPC could plausibly function as intracellular signaling messengers, consistent with the emerging role of lysosomes as signaling organelles.

While prior studies have variably suggested *SPNS1* involvement in lysosomal transport of carbohydrates (*9*) or sphingolipids (*30*), we find that SPNS1 is a pH-dependent lysosomal transporter of lysophospholipids. Notably, our observations are consistent with very recent work by a number of groups, including 1) *He et al.* that described lysophospholipid transport by *SPNS1* (*47*), 2) *Dastvan et al.* that described a proton-driven, alternating-access mechanism for *SPNS1* transporters (*48*); and 3) *Zhou et al.*that predicted a lysolipid-like transport substrate for SPNS1 based on structural modeling of a bacterial homolog (*30*). Moreover, our work provides a notable advance in understanding the physiological role of SPNS1-mediated lysophospholipid transport by demonstrating its critical importance under conditions of nutrient limitation.

Advancing our understanding of *SPNS1* function could ultimately have important implications for human disease. Defects in lysosomal lipid catabolic pathways and lysosomal transporters are known causes of severe, early-onset and age-related neurodegenerative diseases (*29*, *49*)For instance, deficiencies in acid sphingomyelinase (catalyzing lysosomal sphingomyelin hydrolysis), NPC1 and CLN3 (lysosomal lipid or lipid-derived metabolite transporters), cause pathologic accumulation of lysosomal storage materials and present clinically as lysosomal storage diseases with early-onset and fatal neurodegeneration (*29*). To date, *SPNS1* has not been identified as a genetic cause of a neurodegenerative disorder in humans. Nevertheless, deficiency in *SPNS1’s* orthologs has been shown to lead to neurodegeneration in flies (*10*) and to lysosomal dysfunction and storage phenotypes in zebrafish models (*43*). Thus, while germline loss of function *SPNS1* mutations are not compatible with mammalian life, *SPNS1* deficiency in cells and model organisms represents a novel model for lysosomal lipid storage and dysfunction of potential relevance to the study of lysosome-related pathophysiologic mechanisms of neurodegenerative disease.

*SPNS1* has also been identified as a genetic risk factor for various health disorders. Notably, it was found to be a negative prognostic marker in cancer (*50*, *51*). Because choline is highly enriched in rapidly proliferating tumors that require it for lipid biosynthesis, it follows that cancer cells may induce *SPNS1* expression as a cellular strategy to salvage phospholipids for biomass production. *SPNS1* was additionally discovered to be a host factor facilitating SARS-CoV-2 virus entry into lung tissue, possibly through supplying lysophospholipids that enhance membrane fusion (*52*). *SPNS1* is therefore a candidate therapeutic target in cancer and the treatment of viral illnesses. Ultimately, our understanding of the biochemical and physiologic functions of *SPNS1* are still in the nascent stages, and follow-up investigations are needed to characterize the pathophysiologic connections and study the implications of perturbing lysophospholipid efflux through SPNS1 in physiology and disease.

In summary, through uncovering *SPNS1* and its role as an LPC transporter in a novel lysosomal phospholipid salvage pathway, our work demonstrates the power of our platform that combines endolysosomal CRISPR-Cas9 screens with engineered lysosomal dependencies to uncover the biochemical functions of orphan lysosomal proteins and previously undiscovered metabolic pathways.

## Author Contributions

Conceptualization, M.A-R. and S.G.S.; Methodology, S.G.S., W.D., K.N. A.R.K, and M.A-R.; Metabolic Analysis, S.G.S., W.D. and K.N.; Experiments, S.G.S., W.D., K.N. and A.R.K; Library design and CRISPR analysis, R.L-K., S.G.S, K.S., M.A-R. and M.C.B. Manuscript writing, S.G.S. and M.A-R.; Funding Acquisition, M.A-R.

## Acknowledgments

We thank all members of the Abu-Remaileh laboratory for valuable discussions. Additionally, we thank the Metabolomics Knowledge Center (MKC) at Sarafan ChEM-H and the Stanford Neuroscience Microscopy Service at Wu Tsai Neurosciences Institute. This work was supported by grants from the NIH Director’s New Innovator Award Program (1DP2CA271386) and Stanford Cancer Institute to M.A-R., who is a Pew-Stewart Scholar for Cancer Research, supported by The Pew Charitable Trusts and the Alexander and Margaret Stewart Trust. S.G.S and A.K. are supported by the Stanford Medical Scientist Training Program (NIH T32GM007365) and S.G.S. and K.N. are Kolluri Fellows in the Sarafan ChEM-H Chemistry Biology Interface (CBI) Predoctoral Training Program (NIH 5T32GM139791-03). K.N. is additionally supported by the Bio-X Stanford Interdisciplinary Graduate Fellowship affiliated with the Wu Tsai Neurosciences Institute (Bio-X SIGF: Mark and Mary Steven’s Interdisciplinary Graduate Fellow).

## Competing Interests

M.A-R. is a scientific advisory board member of Lycia Therapeutics. All other authors declare no competing interests.

---

**Supplementary Figure 1:**
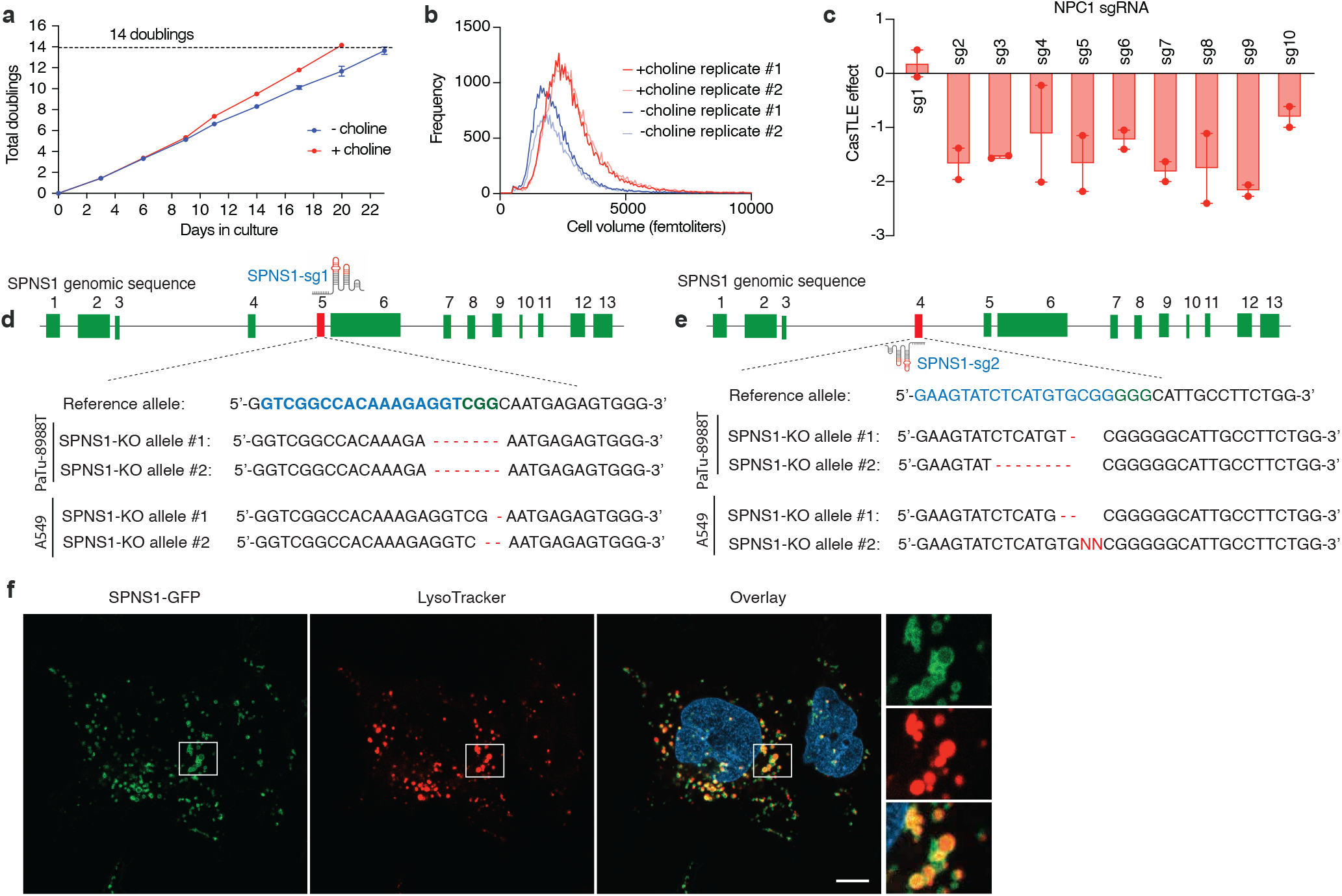
Endolysosomal CRISPR-Cas9 screen implicates SPNS1 In lysosomal choline recycling. **a)** Cell doublings vs. time for PaTu-8988T pancreatic cancer cells grown in choline-depleted medium (-choline, blue) or medium supplemented with 100 μM choline (+choline, red). **b)** Distribution of cell volumes after culturing for twenty days in choline-depleted medium (-choline, blue) or medium supplemented with 100 μM choline (+choline, red). Data in a and b are derived from the actual screen replicates. **c)** Individual CasTLE effects in -choline vs. +choline for all NPC1-targeting sgRNAs used in the endolysosomal library. Data show mean +/− SEM. **d)** Schematic illustrating the genomic target of SPNS1 sgRNA-1 in exon 5 of the human SPNS1 locus and sequences of the mutated alleles of PaTu-8988T-KO1 and A549-KO1 clones. **e)** Schematic illustrating the genomic target of SPNS1 sgRNA-2 in exon 4 of the human SPNS1 locus and sequences of the mutated alleles of PaTu-8988T-KO2 and A549-KO2 clones (bottom). **f)** Fluorescence microscopy images of PaTu-8988T cells expressing SPNS1-GFP. SPNS1-GFP is imaged in the green channel (left). Lysosomes are stained with LysoTracker DND-99 and imaged in the red channel (middle). The merged image (right) demonstrates co-localization of SPNS1-GFP with lysosomes. White scale bar represents 5 μm.

**Supplementary Figure 2:**
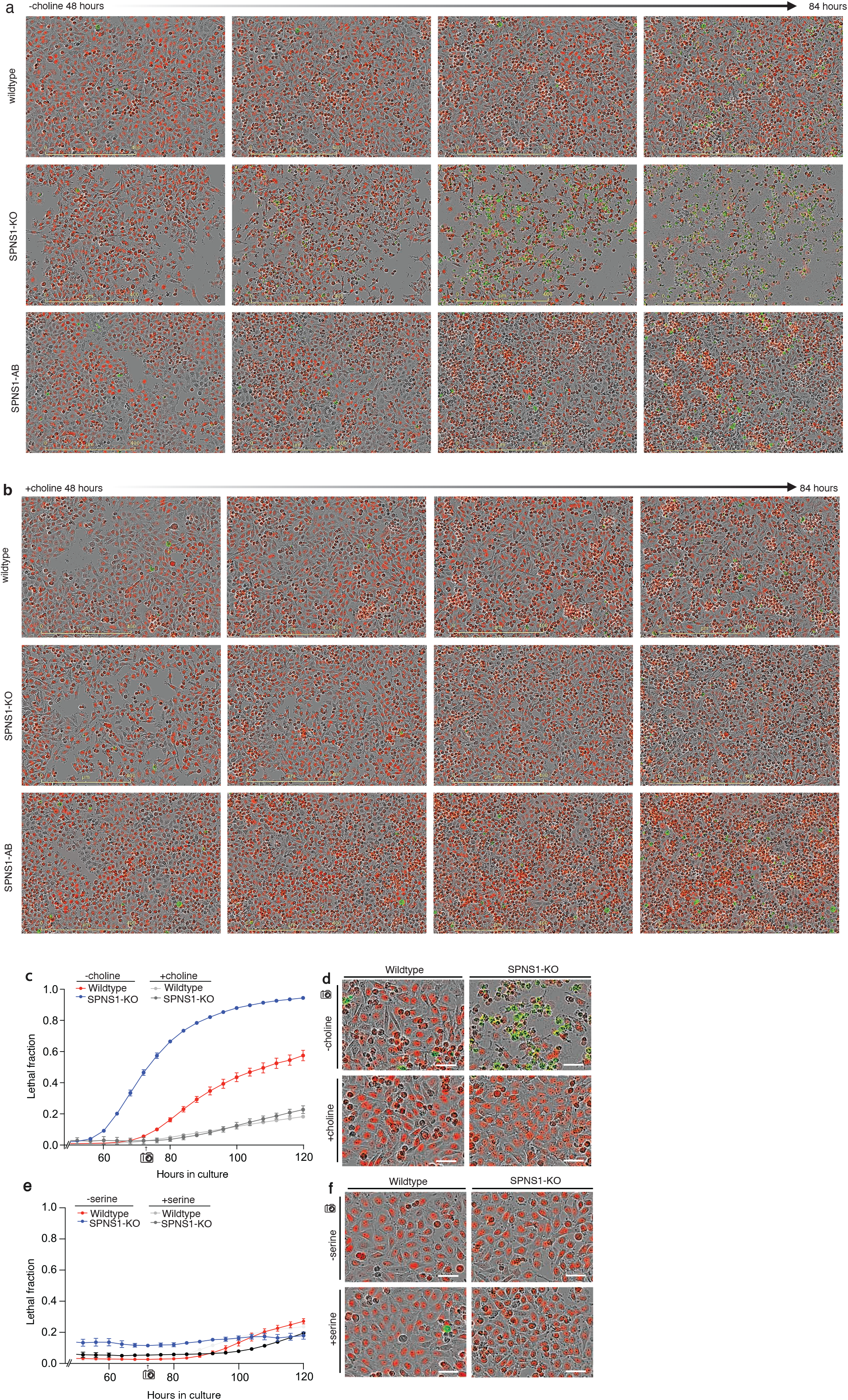
SPNS1 deficiency promotes cell death under choline-limiting conditions. **a&b)** Representative fields from the Incucyte lethal fraction analysis depicting a time-course of wildtype, SPNS1-KO and SPNS1-AB cells during - choline culture (a) and +choline culture (b) from 48 to 84 hours. Green marks dead cells stained with Sytox Green viability dye; red marks nuclei of live cells. From same experiment in Fig. 2c&d. **c)** Lethal fraction of PaTu-8988T wildtype and another SPNS1-KO cell clone grown in choline-depleted medium (-choline) or medium supplemented with 100 μM choline chloride (+choline). Data show mean +/− SEM. **d)** Representative Incucyte fields from (c) at the 72-hour time-point. Green marks dead cells stained with Sytox Green viability dye; red marks nuclei of live cells. **e)** Lethal fraction of wildtype and SPNS1-KO cells (same KO clone as in c) grown in serine-depleted medium (-serine) or medium supplemented with 27 mg/L serine (+serine). Data show mean +/− SEM. **f)** Representative Incucyte fields from (e) at the 72-hour time-point. Green marks dead cells stained with Sytox Green viability dye; red marks nuclei of live cells. White scale bar represents 50 μm.

**Supplementary Figure 3:**
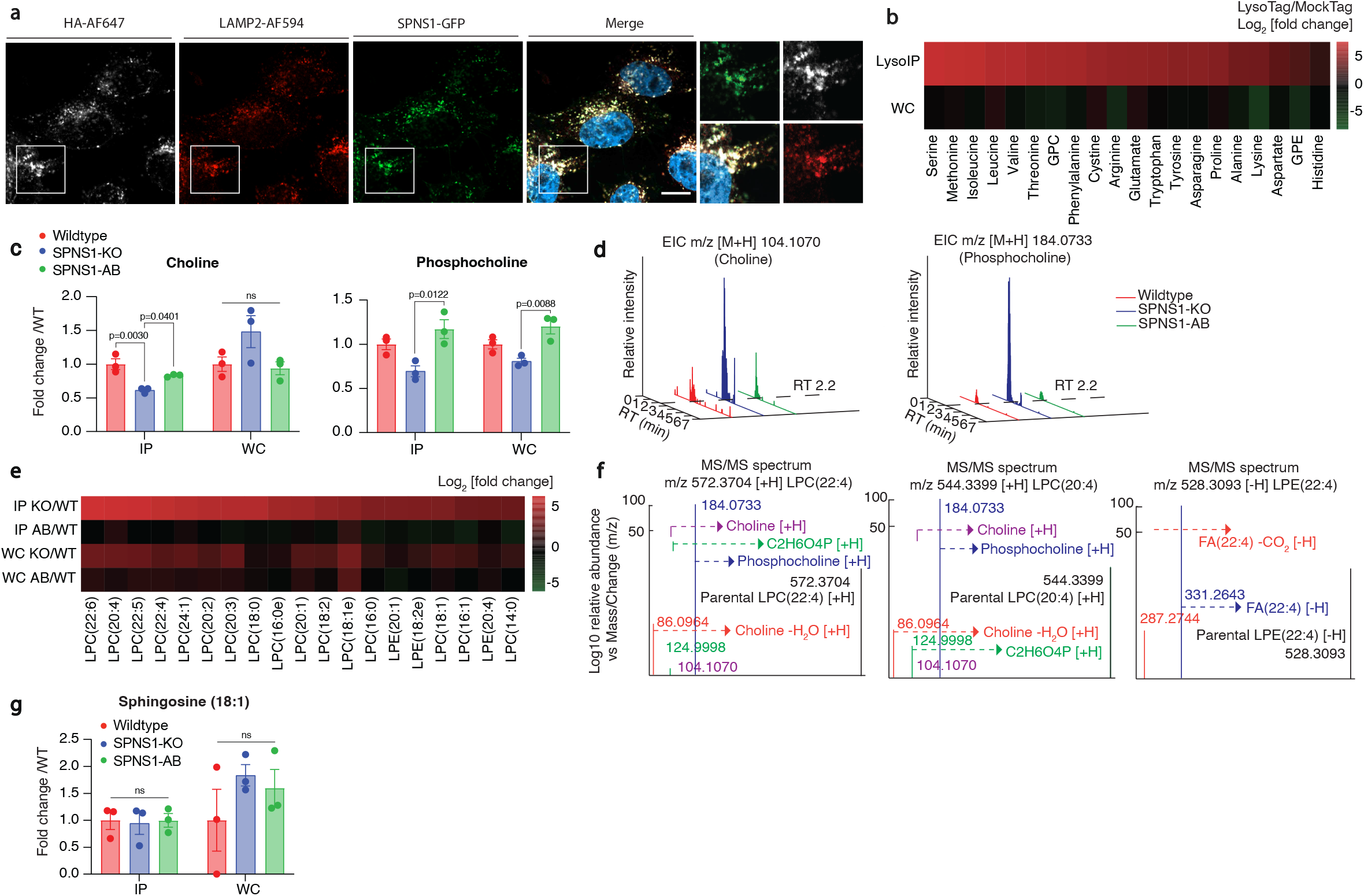
SPNS1-deficient lysosomes accumulate LPCs and LPEs. **a)** Representative fluorescence microscopy images of PaTu-8988T SPNS1-AB cells infected with LysoTag (TMEM192-3xHA). The image depicts HA (LysoTag) staining with AF-647 (white), lysosomal marker LAMP2 staining with AF-598 (red), SPNS1-GFP (green), and a merged image demonstrating co-localization of LysoTag, LAMP2 and SPNS1-GFP. Nuclei are stained with Hoechst (blue). White scale bar represents 5 μm. **b)** Heat map depicting relative abundances of known lysosomal-enriched metabolites in whole-cell (WC) and lysosomes (LysoIP) of wildtype PaTu-8988T cells with LysoTag (TMEM192-3xHA; n=3) versus MockTag (TMEM192-3xflag; n=1). **c)** Quantitation of lysosomal (IP) and whole-cell (WC) abundance of choline and phosphocholine (normalized to a pool of endogenous amino acids: phenylalanine, methionine and tyrosine in positive ion mode) in wildtype, SPNS1-KO1 and SPNS1-AB1 cells. Data show fold change of median of metabolite abundance versus wildtype. Statistical test: one-way ANOVA with Tukey’s HSD post-hoc. **d)** Extracted ion chromatograms (EICs) for choline and phosphocholine produced through in-source fragmentation of lysophosphatidylcholines in wildtype, SPNS1-KO and SPNS1-AB cells. **e)** Heatmap depicting fold changes in abundance of LPC and LPE from hydrophilic interaction liquid chromatography (HILIC, for polar metabolites) measurement in the indicated cells. Within each column (lipid species) for each fraction (IP and WC), fold changes were calculated by normalizing to that of the wildtype (WT) amount.**f)** Annotated tandem mass spectra validating the detection of selected LPCs and LPE. Relative abundance is shown in logarithm scale for better visualization of characteristic fragments with low intensity. **g)** Quantitation of sphingosine (18:1) in wildtype, SPNS1-KO and SPNS1-AB cells demonstrating no accumulation in SPNS1-KO lysosomes (IP) and cells (WC). Data show mean +/− SEM. Statistical test: one-way ANOVA with Tukey’s HSD post-hoc.

**Supplementary Figure 4:**
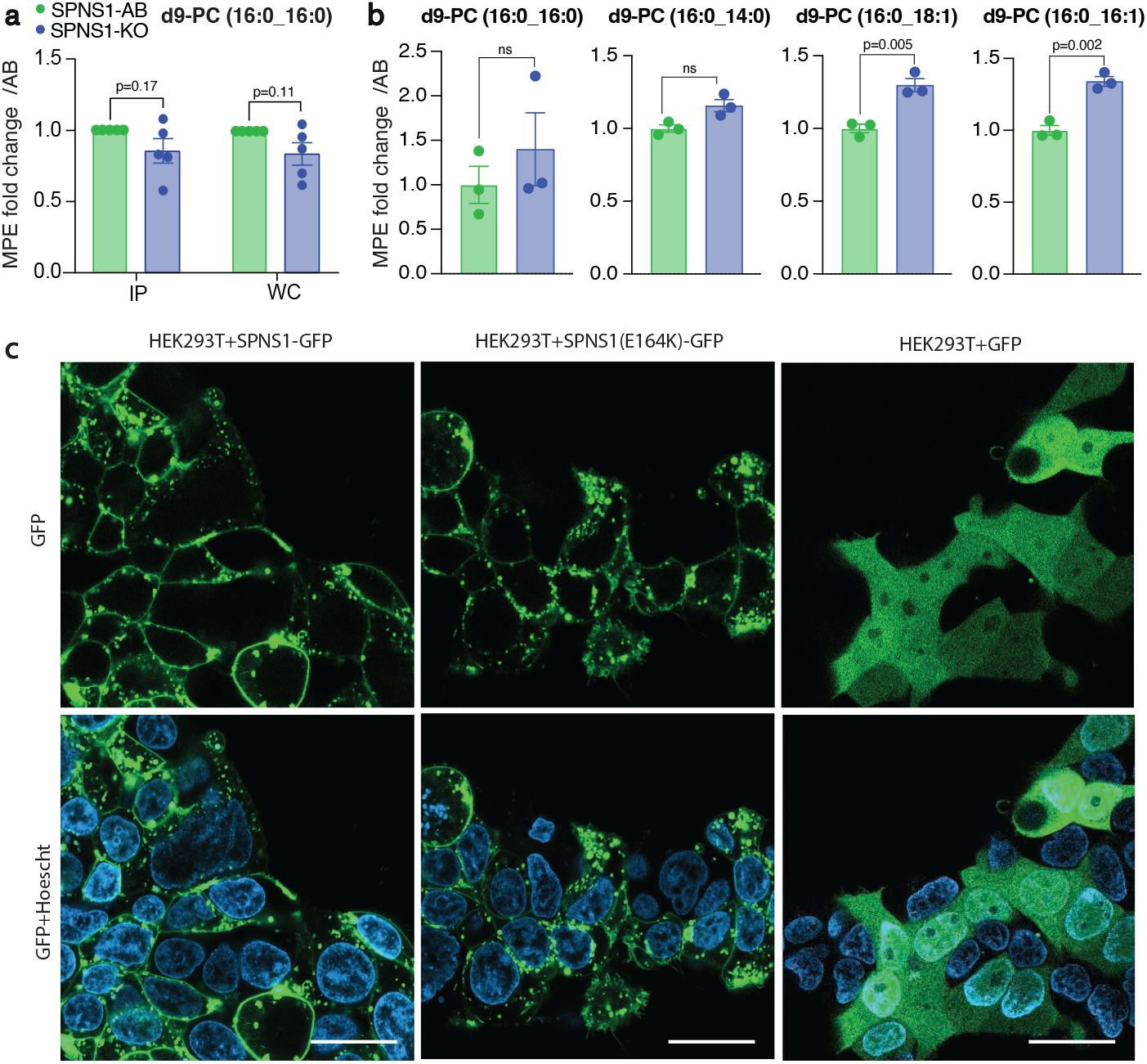
SPNS1 is required for lysosomal efflux of LPC. **a)** Quantitation of tracer d9-PC (16:0_16:0) in lysosomes (IP) and whole cells (WC) of wildtype, SPNS1-KO and SPNS1-AB cells expressed as paired-sample fold-change. Data show mean +/− SEM. Statistical test: two-tailed paired t-test. Data belong to experiments presented in Fig. 4e-f. **b)** Quantitation of molar percent enrichment (MPE) for selected phosphatidylcholines at the whole-cell level. Cells were treated by d9-choline for 3h under choline-replete condition. Data shown as mean +/− SEM. Statistical test: two-tailed unpaired t-test. **c)** Representative fluorescence microscopy images of HEK293T cells over-expressing SPNS1-GFP, SPNS1 (E164K)-GFP and soluble GFP. Nuclei are stained with Hoechst (blue). White scale bar represents 20 μm.

## SUPPLEMENTARY MATERIALS

### Supplementary tables

**Supplementary table 1:** Endolysosomal library sgRNA sequences

**Supplementary table 2:** Ranking of all gene CasTLE effects and scores computed between -choline and +choline conditions in the endolysosomal screen

**Supplementary table 3**: Untargeted lipidomics of the lysosomes and whole cells derived from *SPNS1*-KO and wildtype PaTu-8988T cells

### Materials and Methods

#### Cell culture

Unless otherwise indicated, all cell lines and their derivatives were cultured in DMEM (Gibco) supplemented with 10% heat-inactivated fetal calf serum (FBS) (Thermo Fisher Scientific), GlutaMAX™ (Gibco), and penicillin and streptomycin (Thermo Fisher Scientific). All cell lines were maintained at 37°C and 5% CO2. All cell lines were frequently tested for mycoplasma.

#### Recombinant virus production and transduction

##### Production

Viral HEK293T cells were grown to 40-60% confluency and co-transfected with lentiviral or retroviral packaging plasmid, VSV-G pseudotyping plasmid, and vector (or vector library) plasmid at a mass ratio of 9:1.5:10, respectively, using the XtremeGene 9 (Roche) transfection reagent. 16 hours post-transfection, culture media was replaced with DMEM media supplemented with 30% FBS and penicillin/streptomycin. The virus-containing supernatant was harvested after 48 hours (first collection) and 96 hours (second collection), centrifuged to remove cell debris, and filtered through a 0.22 μM syringe filter.

##### Transduction

Two million cells were plated in six-well plates in DMEM with the addition of 8μg/mL polybrene and 100–250μl of virus-containing medium. Spin-infection was performed at 2,200 rpm for 45 minutes at 37 °C, and cells were incubated with virus for 16 hours before adding fresh culture medium containing the relevant antibiotic for selection for at least 72 hours.

#### Endolysosomal CRISPR-Cas9 screen

##### Endolysosomal library construction

A list of lysosomal, endocytic and autophagy-related genes in addition to a selective set of metabolism and signaling genes (1,061 genes) was manually curated by unbiased profiling of lysosomal proteins using in-house LysoIP proteomics and by selecting genes implicated in endolysosomal trafficking (*53*). For the library design, 10 targeting sgRNAs were selected per gene, and an additional 1,050 sgRNAs (~9% of the library) were included as safe-targeting (control) sgRNAs. These guides were cloned into pMCB320 as previously described (*54*).

##### Endolysosomal screen

The endolysosomal library was infected into Cas9-expressing PaTu-8988T pancreatic cancer cells at 1000x library coverage and selected using puromycin. At time point 0, cells were trypsinized, washed, and split into two culture media conditions: choline-depleted medium (-choline) or choline-supplemented medium (+choline). Choline-depleted medium was comprised of 1:1 DMEM/Ham’s F12 without sodium bicarbonate, serine, methionine, or choline chloride (Caisson Labs; DFP14-5) supplemented with 1200 mg/L sodium bicarbonate, 18 mg/L methionine, 27 mg/L serine, 5% triple-dialyzed FBS and penicillin/streptomycin. To make +choline medium, -choline medium was supplemented with 100 μM choline chloride.

Cells were propagated in -choline or +choline medium for 14 doublings while maintaining a library coverage of >1000x. DNA from time point 0 (T0) and after fourteen doublings for each condition was isolated and sgRNA sequences were PCR-amplified. Illumina universal adaptor sequences were subsequently added to sgRNA amplicons by PCR, and sgRNA abundances were determined using the Illumina NextSeq 550 with NextSeq 500/550 mid-output kit v2.5.

##### Screen analysis

Sequencing data were analyzed using the CasTLE method developed by *Morgens et al. (17*). In brief, the method calculates the most likely maximum effect (phenotype) size (CasTLE effect) among each group of gene-targeting sgRNAs by comparing each set to the negative sgRNAs (control and safe-targeting) in the library. The method then scores the significance of the effect (CasTLE score) by permuting the results.

#### Generation of knockout cell lines

The following *SPNS1* sgRNAs were used in this study: *SPNS1*-sg1: 5’-GTCGGCCACAAAGAGGT-3’; *SPNS1*-sg2: 5’-GAAGTATCTCATGTGCGG-3’. *SPNS1*-targeting sgRNAs were cloned into the Px458 vector containing GFP as a selectable marker. sgRNA-Px458 plasmids were transfected into cells using XtremeGene 9 transfection reagent and 48 hours post-transfection GFP+ cells were single-sorted into 96 well plates prepared with 200 μL/well DMEM with 30% FBS and penicillin/streptomycin. Plates were incubated at 37 °C and 5% CO2 for three weeks and routinely checked for colony formation. Colonies were harvested and sgRNA target sites PCR-amplified and sanger-sequenced. Indels generated at the sgRNA target site were determined using Synthego ICE indel deconvolution web-based tool (https://www.synthego.com). Cathepsin B, D and L knockout lines were generated using pLentiCRISPRv1 and previously validated (*55*).

#### Screen hits validation by endpoint viability

Screen hits were validated by generating knockout cell lines and comparing viability between cells cultured in -choline and +choline medium after five days. Cells were seeded in triplicate in 96-well opaque-edge clear-bottom plates (Corning) at a density of 2000 cells/well and incubated at 37 °C and 5% CO2 until the most confluent wells reached ~95% confluence (approximately five days) without media change. Cell viability was measured using the CellTiter-Glo luminescent viability assay (Promega) on a SpectraMax i3 plate reader (Molecular Devices).

#### Incucyte live cell analysis and lethal fraction calculations

Lethal fraction curves were produced using an Incucyte S3 (Sartorious) as previously described (*31*). Briefly, live cells were labeled with nuclear-localized red fluorescent protein mKate2 and the frequency of live cells determined by counting mKate2+ (red) nuclei per field; dead cells were labeled using SYTOX green (SG) viability dye (Invitrogen) and the frequency of dead cells determined by counting green cells per field. mKate2^+^and SG^+^ (double-positive) cells detected in the overlap channel were defined as dead cells (recently dead cells may stain double-positive due to residual mKate2 signal). Lethal fraction was calculated as 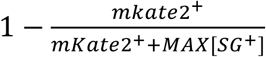, where mKate2^+^ cells excludes double-positive mKate2^+^SG^+^ cells, and MAX[SG^+^] is the maximum number of SG^+^ cells observed in the given field at any analysis timepoint (the MAX[SG^+^]metric accounts for the disappearance of dead cells over time). Incucyte analysis parameters were optimized for the detection of live and dead cells for the PaTu-8988T cell type. Parameters are tabulated below and were kept consistent across all lethal fraction analyses.

| **Green parameters** | **Selection** | **Specification** |
| --- | --- | --- |
| Segmentation | Adaptive | Threshold: 1.5 GCU |
| Edge sensitivity | Edge split: on | Sensitivity: -30 |
| Filters | Area | Min: 50 µm; Max: 750 µm |
| **Green parameters** | **Selection** | **Specification** |
| Segmentation | Top-Hat | Radius: 100; Threshold: 3.0 RCU |
| Edge sensitivity | Edge split: on | Sensitivity: -10 |
| Filters | Area | Min: 50 µm; Max: 1000 µm |

For cell lines not infected with mKate2 red fluorescent nuclear protein, cell death was assessed using Propidium Iodide viability stain. The Incucyte system was used to image PI staining over time or at one end-point.

#### Cell growth curves

Cell growth curves were generated by assessing live-cell abundance using CellTiter-Glo Luminescent Cell Viability Assay (Promega). Cells were seeded in triplicate at a density of 2000 cells/well in seven separate opaque-walled plates. One plate was used for viable cell analysis by CellTiter-Glo at each timepoint to generate a time-dependent growth curve.

#### Immunoblotting

Cell lysates were resolved by SDS-PAGE (Thermo Fisher Scientific) at 80 V and transferred to PVDF membranes for 2 hours at 40 V. Membranes were blocked with 5% BSA (bovine serum albumin) in TBST buffer (Tris-buffered saline with Tween-20) for 30 minutes, then incubated overnight with primary antibodies in 5% BSA in TBST at 4 °C. The following primary antibody dilutions were used: LAMP2 (H4B4, Santa Cruz Biotechnology, 1:1000); VDAC (B-6, Santa Cruz Biotechnology, 1:1000); Cathepsin B (D1C7Y, Cell Signaling, 1:1000). After incubation, membranes were washed three times with TBST for 15 minutes per wash and incubated with the appropriate secondary antibodies diluted 1:3,000 in 5% BSA in TBST for 1 hour at room temperature. Finally, membranes were washed three times with TBST and visualized using ECL2 western blotting substrate (Thermo Fisher Scientific) on a ChemiDoc MP imaging system (BioRad).

#### Immunofluorescence

Cells plated on glass coverslips were fixed with 4% paraformaldehyde and blocked with 3%BSA in PBS for 1 hour at room temperature. Primary antibodies were incubated overnight at 4°C with dilutions as follows: mouse anti-LAMP2 (1:500), rabbit anti-HA (1:500). Secondary antibodies (Alexa-Fluor) were then applied in 1% BSA, 0.3% Triton X-100 for 3 hours. The following secondary antibodies were used (1:1000): goat antimouse Alexa Fluor 647 (Thermo Fisher Scientific, A-21240), and goat anti-rabbit Alexa Fluor 594 (Thermo Fisher Scientific, A32740). Confocal images were acquired on the Zeiss LSM980 with Airyscan 2.

#### Cell volume measurement

Cell volumes were determined using a Beckman Z2 particle counter and size analyzer with filtering criteria set to include cells between 10 μm and 30 μm.

#### LysoIP

Lysosome purification by LysoIP was adapted from the method previous described (*4*) and optimized for the PaTu-8988T cell line. Because we observed better enrichment post-optimization in *SPNS1*-KO clone #2 compared to clone #1, we used #2 for all main figure LysoIP and tracing experiments. Similar LysoIP results obtained with the other clone. Briefly, cells were grown to 95% confluence in 15 cm cell culture dishes. If lysosomal isolates were to be used for downstream metabolomics analysis, 4 μM Lysotracker Red DND-99 (Invitrogen) was added to each plate 1 hour prior to cell harvest. At the time of harvest, each plate was washed 2x with 5 mL of ice-cold PBS and cells were scraped in 1 mL ice cold KPBS (136 mM KCl, 10 mM KH_2_PO_4_, pH 7.25 in Optima LC–MS water) and transferred to a 2 mL tube. Cells were pelleted by centrifugation at 1000*xg* for 2 minutes at 4 °C. Following centrifugation, cells were resuspended in 500 μL fresh KPBS and 12.5 μL of the cell suspension was taken for whole-cell fraction analysis. The remaining cell suspension was lysed by trituration using a 29.5-gauge insulin syringe (EXELINT international), then diluted with an additional 500 μL KPBS and centrifuged at 1000*xg* for 2 minutes at 4°C to pellet cell debris. The lysosome-containing supernatant (1mL) was added to tubes containing an equivalent of 100 μL Pierce anti-HA magnetic beads (Thermo Scientific) and rocked for 3 minutes at 4 °C. Following incubation, the bead-lysosomes complexes were washed 3x with ice cold KPBS and extracted in the appropriate buffer. For metabolite extraction, lysosome and whole-cell fractions were extracted in 50 μL and 225 μL, respectively, of 80% methanol in LC–MS water containing 500 nM isotope-labeled amino acids used as internal standards (Cambridge Isotope Laboratories). Following extraction, HA-binding beads were removed from the lysosome fractions using a rotary magnet and cell debris was removed from whole-cell extractions by centrifugation at 20,000*xg* at 4 °C for 15 minutes. For lipidomic extraction, lysosome and whole-cell fractions were extracted in 1 ml chloroform:methanol at ratio of 2:1 (v/v) containing 750 ng ml^-1^ of SPLASH LIPIDOMIX internal standard mix (Avanti) for > 10 minutes. Following extraction, HA-binding beads were removed using a rotary magnet and LysoIP and whole-cell samples were vortexed for 1 hour at 4°C. 200 μL saline was then added to each sample, and samples were vortexed for an additional 10 minutes. Vortexed samples were spun at 3000xg for 5 min to separate polar (top) and nonpolar (bottom) phases. 600 μL of the lipid-containing chloroform (nonpolar) phase was dried and reconstituted in Acetonitrile:isopropanol:water 13:6:1 (v/v/v). Both metabolomic and lipidomic extractions were analyzed on mass spectrometer with workflow and parameters described below.

#### Untargeted Metabolomics

Profiling of polar metabolites was performed on an ID-X tribrid mass spectrometer (Thermo Fisher Scientific) with an electrospray ionization (ESI) probe. A SeQuant^®^ ZIC^®^-pHILIC 150 x 2.1 mm column (Millipore Sigma 1504600001) coupled with a 20 x 2.1 mm (Millipore Sigma 1504380001) guard was used to carry out hydrophilic interaction chromatography (HILIC) for metabolite separation prior to mass spectrometry. Mobile phases: A, 20 mM ammonium carbonate and 0.1% ammonium hydroxide dissolved in 100% LC/MS grade water; B, 100% LC/MS grade acetonitrile. Chromatographic gradient: linear decrease from 80-20% B from 0-20 minutes; fast linear increase from 20-80% B from 20-20.5 minutes; 80% B hold from 20.5-29.5 minutes. Flow rate, 0.15 ml/minute. Injection volume, 1.5-2.5 μL. Mass spectrometer parameters: ion transfer tube temperature, 275 °C; vaporizer temperature, 350 °C; Orbitrap resolution, 120,000; RF lens, 40%; maximum injection time, 80 ms; AGC target, 1×106; positive ion voltage, 3000 V; negative ion voltage, 2500 V; Aux gas, 15 units; sheath gas, 40 units; sweep gas, 1 unit. Full scan mode with polarity switching at m/z 70-1000 was performed. EASYIC^™^ was used for internal calibration. For data-dependent MS2 collection, pooled samples were prepared by combining replicates. HCD collision energies, 15, 30 and 45%; AGC target, 2×10^6^; Orbitrap resolution, 240,000; maximum injection time, 100 ms; isolation window, 1 m/z; intensity threshold, 2×10^4^; exclusion duration, 5 seconds; isotope exclusion, enable. Background exclusion was performed via AcquireX with one header blank and the exclusion override factor set to 3.

Rigorous quantification of metabolite abundance was performed by TraceFinder (Thermo Fisher Scientific) in conjunction with an in-house library of known metabolite standards (MSMLS, Sigma-Aldrich). Mass tolerance for extracting ion chromatograms: 5 ppm.

#### Untargeted lipidomics workflow

Profiling of nonpolar lipids was performed on an ID-X Tribrid mass spectrometer (Thermo Fisher Scientific) with a heated electrospray ionization (HESI) probe. An Ascentis Express C18 150 × 2.1 mm column (Millipore Sigma 53825-U) coupled with a 5 × 2.1 mm guard (Sigma-Aldrich 53500-U) was used to carry out C18-based lipid separation prior to mass spectrometry. Mobile phases: A, 10 mM ammonium formate and 0.1% formic acid dissolved in 60% and 40% LC/MS grade water and acetonitrile, respectively; B, 10 mM ammonium formate and 0.1% formic acid dissolved in 90% and 10%LC/MS grade 2-propanol and acetonitrile, respectively. Chromatographic gradient: isocratic elution at 32% B from 0-1.5 minutes; linear increase from 32-45% B from 1.5-4 minutes; linear increase from 45-52% B from 4-5 minutes; linear increase from 52-58% B from 5-8 minutes; linear increase from 58-66% B from 8-11 minutes; linear increase from 66-70% B from 11-14 minutes; linear increase from 70-75% B from 14-18 minutes; linear increase from 75-97% B from 18-21 minutes; hold at 97% B from 21-35 minutes; linear decrease from 97-32% B from 35–35.1 minutes; hold at 32% B from 35.1-40 minutes. Flow rate, 0.26 ml/minutes. Injection volume, 2-4 μL. Column temperature, 55°C. Mass spectrometer parameters: ion transfer tube temperature, 300 ¤C; vaporizer temperature, 375 °C; Orbitrap resolution MS1, 120,000, MS2, 30,000; RF lens, 40%; maximum injection time MS1, 50 ms, MS2, 54 ms; AGC target MS1, 4×10^5^, MS2, 5×10^4^; positive ion voltage, 3250 V; negative ion voltage, 3000 V; Aux gas, 10 units; sheath gas, 40 units; sweep gas, 1 unit. HCD fragmentation, stepped 15%, 25%, 35%; data-dependent tandem mass spectrometry (ddMS2) cycle time, 1.5 s; isolation window, 1 m/z; microscans, 1 unit; intensity threshold, 1.0e4; dynamic exclusion time, 2.5 s; isotope exclusion, enable. Full scan mode with ddMS2 at m/z 250-1500 was performed. EASYIC^™^ was used for internal calibration. LipidSearch and Compound Discoverer (Thermo Fisher Scientific) were used for unbiased differential analysis. Lipid annotation was acquired from LipidSearch with the precursor tolerance at 5 ppm and product tolerance at 8 ppm. The mass list was then exported and used in Compound Discoverer for improved alignment and quantitation. Mass tolerance, 10 ppm; minimum and maximum precursor mass, 0-5,000 Da; retention time limit, 0.1-30 min; Peak filter signal to noise ratio, 1.5; retention time alignment maximum shift, 1 min; minimum peak intensity, 10,000; compound detection signal to noise ratio, 3. Isotope and adduct settings were kept at default values. Gap filling and background filtering were performed by default settings. The MassList Search was customized with 5 ppm mass tolerance and 1 minute retention time tolerance. Area normalization was performed by constant median after blank exclusion.

#### ^2^H-isotope tracing by d9-dipalmitoylphosphatidylcholine (16:0–16:0) in PaTu-8988T cells

*SPNS1*-KO and *SPNS1-AB* PaTu-8988T cells were seeded according to LysoIP procedure. Upon ~95% confluency, the cells were washed by PBS and the medium was replaced with FBS-depleted media for 3 hours for serum starvation. The cells were subsequently labelled with 37.6 μM d9-dipalmitoylphosphatidylcholine (16:0_16:0) conjugated with 0.25% fatty-acid free bovine serum albumin for 2 hours. Whole-cell and lysosomal fractions were obtained according to the LysoIP procedure.

To assess lysosomal contribution to choline-containing lipids at the whole-cell level, molar percent enrichment (MPE) for trimethyl-d9 labeling was calculated based on the following equation:

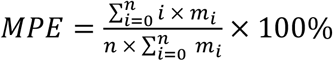

where *n* is the number of hydrogen atoms in the lipid molecule, *mi* the abundance of a mass isotopomer and *i* the labeling state (M+i). As a choline-containing lipid can only be present as either unlabeled (M+0) or d9-labeled (M+9), the equation above is simplified as follows:

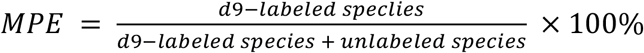

#### ^2^H-isotope tracing by d9-choline in PaTu-8988T cells

*SPNS1*-KO and AB PaTu-8988T cells were seeded at 1.5 million per well in 6-well plates. Upon full confluency, the cells were first washed by PBS and then labeled with 37.6 μM d9-choline chloride mixed with 0.25% fatty-acid free bovine serum albumin for 3 hours. The whole-cell harvest was then processed for total lipid analysis. MPE was calculated as mentioned previously.

#### Transport assay

Equal numbers of wildtype and *SPNS1*-overexpressing HEK293T cells were incubated in 100 μL transport assay buffer with the indicated concentration of d7-LPC (18:1) (Avanti) for 20 minutes at 37 °C. The composition of transport assay buffers was as follows: pH 5: 25 mM sodium acetate, 5 mM glucose, 1 mM MgCl, 150 mM NaCl; pH 7: 25 mM HEPES, 5 mM glucose, 1 mM MgCl, 150 mM NaCl. Following incubation, cells were placed on ice and immediately washed twice with 0.5% fatty acid-free BSA in PBS and once with PBS. Lipids were extracted in acetonitrile:isopropanol:water 13:6:1 (v/v/v) for 15 minutes and spun at 20,000xg for 10 minutes at 4 °C to remove cell debris. Lipid extracts were analyzed on the Agilent Triple-Quadrupole 6470 Mass Spectrometer as described below.

#### d7-LPC (18:1) quantitation

Lipids were separated on an Ascentis C18 column (5 Micron, 5 mum particle size, L x I.D. 5cm x 4,6mm) (Sigma-Aldrich) with an Ascentis Express guard holder and connected to a 1290 LC system. The LC system was coupled to the 6470A triple quadrupole (QQQ) mass spectrometer equipped with an LC-ESI probe. External mass calibration was performed using the standard calibration mixture every 7 days. Injection volumes of 2-4 μL were used for each sample, with separate injections for positive and negative ionization modes. Mobile phase A in the chromatographic method consisted of 60:40 water:acetylnitrile with 10mM ammonium formate and 0.1% formic acid, and mobile phase B consisted of 90:10 isopropanol:acetylnitrile, with 10 mM ammonium formate and 0.1% formic acid. The chromatographic gradient was adapted from Laqtom *et al. (38*). In brief, the elution was performed with a gradient of 18 min; during 0-1 min isocratic elution with 32% B, increase to 66% B from 1 to 6 min, increase to 75% B; from 6 to 10 min, increase to 97% B; from 10 to 14 min, solvent B was decreased to 32% and then maintained for another 4 min for column re-equilibration. The flow rate was set to 0.260 mL/min. The column compressor and autosampler were held at 55 °C and 4 °C, respectively. The mass spectrometer parameters were as follows: the spray voltage was set to 3.5 kV in positive mode and 2.5 kV in negative mode, and the gas temperature and the sheath gas flow were held at 250 °C and 300 °C, respectively. Both gas flow and sheath gas flow were 12 L/min while the nebulizer was maintained at 25 psi. These conditions were held constant for both positive- and negative-ionization mode acquisitions.

The mass spectrometer was operated in Multiple Reaction Method (MRM) for targeted analysis of species of interest. Standard compounds including d7-LPC (18:1), d7-LPE (18:1), d5-LPC (15:0), LPC (18:1), LPC (15:0), PI (16:0_18:1) and PC(16:0_18:0) (POPC) were purchased from Avanti Polar Lipids and optimized using a MassHunter Optimizer MRM. MassHunter Optimizer MRM is an automated method development software used to generate and optimize MRM transitions accumulating at most 4 products with different abundances from singly ionized species. The two most abundant transitions from either the negative or positive mode were selected to detect each species. The precursor-product ion pairs (m/z) used for MRM of the compounds were as follows: d7-LPC 18:1: 529.4→104.1/184.0; d7-LPE 18:1: 487.4→346.3/62.1; d5-LPC 15:0: 487.4→184.0/104.1; LPC 18:1: 522.4→281.3/104.1; LPC 15:0: 482.4→60.2/104.1; PI 16:0/18:1: 854.6→577.5/837.5 and POPC: 760.6→124.9/60.2.

High-throughput annotation and relative quantification of lipids were performed using a qualitative analysis software of MassHunter acquisition data and QQQ quantitative analysis (Quant-My-Way) software. Individual lipid species shown in the figures were validated using the Qualitative software by manually checking the peak alignment and matching the retention times and MS/MS spectra to the characteristic fragmentation compared to the standard compounds. Analyzing two transitions for the same compound and looking for similar relative response was an added validation criterion to ensure the correct species were identified and quantified. The MRM method and retention time were used to quantify all lipid species using the quantification software, and the raw peak areas of all species were exported to Microsoft Excel for further analysis. d7-LPC raw abundances were normalized to cell number using abundance of endogenous control lipids in the same sample.

#### Multiple sequence alignment

Sequences of Homo sapiens SPNS1 (Q9H2V7) and Drosophila melanogaster Spinster (alias: bnch, Q9GQQ0) were obtained from UniProt. Clustal Omega was used to perform a multiple sequence alignment using inputted protein sequences.

#### Docking with Schrodinger Glide

I-TASSER (*56*) was used to predict the structure of Homo sapiens SPNS1 using Q9H2V7 (UniProt) as the entry sequence. The predicted Homo sapiens SPNS1 structure was prepared in Schrödinger Glide using the Protein Preparation Wizard, which ensures structural correctness of the molecule, for example by filling in missing residues, flipping positions of residues, adding H atoms, adding appropriate bond orders, and adding in formal charges to ensure that all atoms and residues are represented accurately in the protein. The structure of LPC 22:4 was obtained from the ZINC15 database (ZINC000040165293) and prepared using Schrödinger’s LigPrep tool to generate a 3D ligand structure with energy minimization. Afterward, Schrödinger’s Receptor Grid Generation tool was used to create a grid representing the shape and properties of the receptor binding site for use in docking simulations using the centroid of three residues around the putative substrate binding site. Finally, using the prepared ligand library and receptor grids, Glide Ligand Docking was performed using standard precision, which accounts for hydrophobic interactions, hydrogen bonding, and coulombic, van der Waals, and solvation terms.

#### Data preparation and statistics

Displays of quantitative data were prepared in Microsoft Excel v.16.65 and GraphPad Prism v.9.0. Statistical comparisons were performed using two-tailed unpaired or paired *t-*tests and ordinary one-way ANOVA in Prism or Excel, unless stated otherwise in the figure legends. All displayed measurements represent samples generated independently or biological replicates unless otherwise indicated. Figure schematics were created using BioRender and were used with permission.

